# Predicting the Prognosis of Non-Small Cell Lung Cancer by Integrating Microarray and Clinical Data with Deep Learning

**DOI:** 10.1101/656140

**Authors:** Yu-Heng Lai, Wei-Ning Chen, Te-Cheng Hsu, Che Lin, Yu Tsao, Semon Wu

## Abstract

Non-small cell lung cancer (NSCLC) is one of the most common lung cancers worldwide. Accurate prognostic stratification of NSCLC can become an important clinical reference when designing therapeutic strategies for cancer patients. With this clinical application in mind, we developed a deep neural network (DNN) combining heterogeneous data sources of gene expression and clinical data to accurately predict the prognosis of NSCLC patients. Based on microarray data from a cohort set (614 patients), seven well-known NSCLC markers were used to group patients into marker- and marker+ subgroups. Using a systems biology approach, prognosis relevance values (PRV) were then calculated to select eight additional novel prognostic gene markers. Gene markers along with clinical data were then used to develop an integrative DNN via bimodal learning to predict the 5-year survival rate of NSCLC patients with tremendously high accuracy (AUC: 0.8163, accuracy: 75.44%), which is superior to all other existing methods based on AUC. Using the capability of deep learning, we believe that our predicted cancer prognosis can be a promising index helping oncologists and physicians develop personalized therapy and build the foundation of precision medicine in the future.

## 1 Introduction

Lung cancer is the worldwide leading cause of cancer-related mortality, with non-small cell lung cancer (NSCLC) accounting for approximately 85% of all lung cancer patients [1]. The most common NSCLC subtypes are adenocarcinoma (ADC), squamous cell carcinoma (SQC), and large cell carcinoma. Although the overall 5-year survival rate of patients diagnosed with stage I ADC was 63%, nearly 35% of patients relapsed after surgery with a poor prognosis [2]. Adjuvant treatments have been considered ideal for ADC patients with the highest risk of recurrence or death to increase survival rates [3]. Therefore, prognostic stratification is crucial for categorizing patients to help doctors make decisions on therapeutic strategies.

In recent years, researchers have developed predictive methods based on gene expression profiles to classify lung cancer patients with distinct clinical outcomes, including relapse and overall survival [4]. Previous studies have shown the importance of biomarkers for NSCLC, such as EPCAM, HIF1A, PKM, PTK7, ALCAM, CADM1, and SLC2A1, which were used as a single biomarker for predicting prognosis or metastasis [5, 6, 7, 8, 9, 10, 11]. However, cancer is a systemic disease with complicated and illusive mechanisms that often involves multiple genes and cross-talk between pathways. Therefore, extending our understanding of NSCLC via the single gene biomarkers by studying the interactions between genes is essential for more accurate prognosis prediction.

Machine learning algorithms are powerful tools that apply input features (biomarkers) to capture the complicated interdependencies between these features to accurately predict clinical outcomes based on the considered dataset [12]. In addition, predicting cancer prognosis can be improved by appropriately modeling the interactions between biomarkers compared with the single biomarker approach [13].

Due to the relative small sample size of patient data compared to the large number of genetic features, scientists have assiduously focused on feature selection algorithms that aim to obtain a subset of significantly representative features [14]. However, while traditional feature selection methods are often based on the statistical or predictive performance of the patient data set, biological concepts were rarely considered when isolating potential gene features. Therefore, the predicted gene features to apply and improve therapeutic strategies for cancer patients are limited.

Systems biology is computational and mathematical modeling of complex biological systems that has been widely applied [15]. There is increasing interest in applying systems biology approaches to identify cancer-associated genes as feature selection strategies [16]. In this study, we established a systems biology approach for NSCLC patients, which identified eight novel survival-related genes based on seven previously well-known markers. Among these prognostic markers, selected features were used in the following machine learning models to predict the survival rate of patients.

Deep learning has seen unprecedented success in many fields, such as image recognition [17], speech recognition [18], and biology [19]. A deep neural network (DNN) is composed of non-linear modules, which represent multiple levels of abstraction [20]. Each representation can be transformed into a slightly more abstract level, leading to even more involved interactions among features. As a result, deep learning algorithms can extract high-level abstractions from different types of data sources and provide superior performance compared with traditional machine learning methods. Thanks to the representation of features in hidden layers, a DNN can easily combine the networks for different modalities. As a result, we aimed to propose an integrative DNN that combines both gene expression and clinical data to improve prognosis prediction for NSCLC patients.

In this study, we integrated a systems biology approach that combined seven well-known biomarkers and identified eight biologically significant prognostic gene features with a deep learning method. We combined gene expression and clinical data sources to predict the 5-year survival of NSCLC patients. We believe our significant improvements to predicting prognostic outcomes for lung cancer patients may help oncologists and physicians make accurate and precise decisions on appropriate treatment for individual patients, which may build the foundation for future personalized therapeutic strategies.

## 2 Results

We integrated systems biology and deep learning approaches to predict the survival status of NSCLC patients. In addition, the systems biology approach was specifically used to identify prognostic markers (gene features). The selected prognostic markers were used as input features for our DNN prediction based on the 5-year survival of lung cancer patients. Moreover, we further integrated their clinical background via an integrative DNN model to improve the performance of our prediction. The schematic of our strategy is shown in Figure. 1.

**Figure 1:**
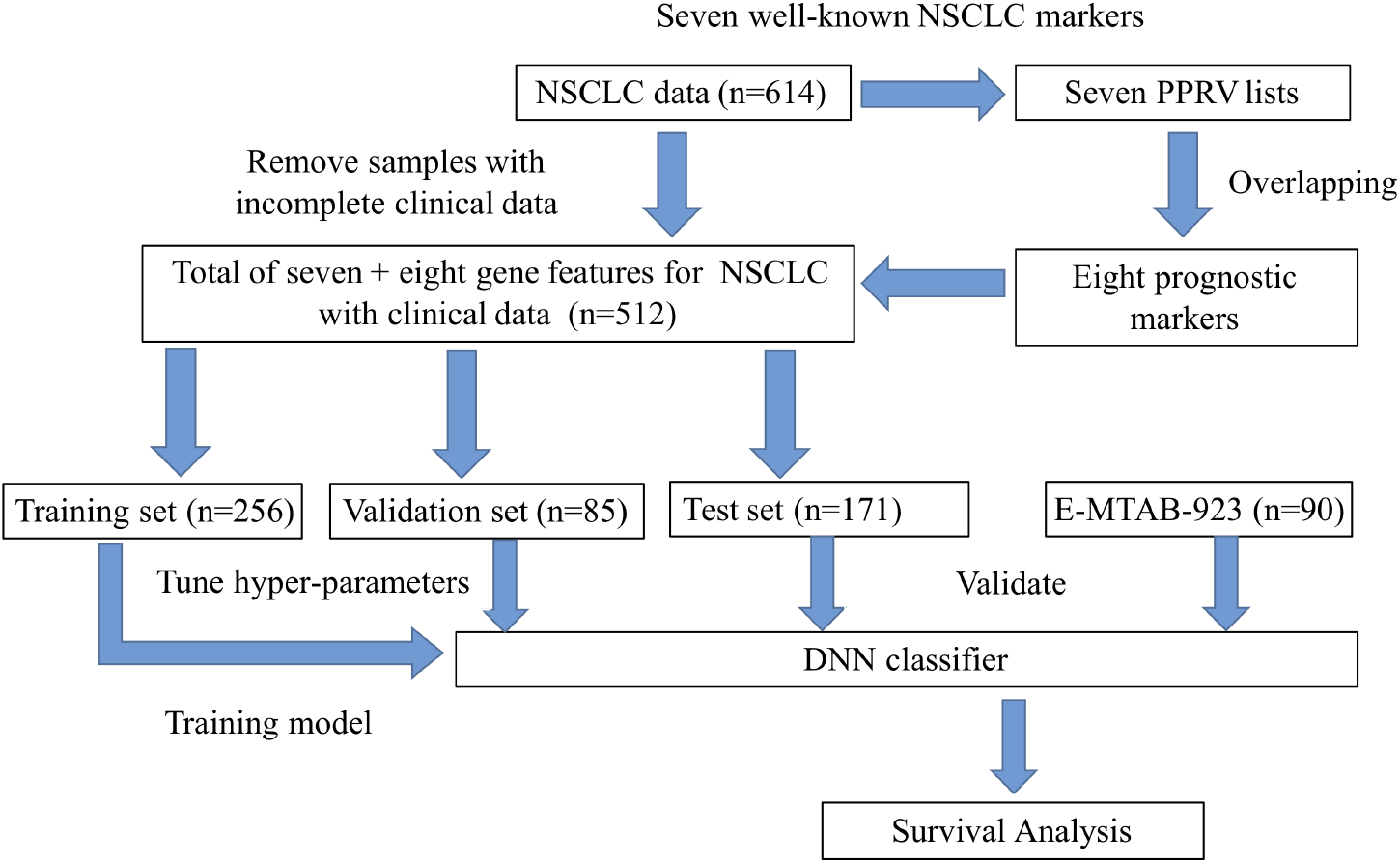
Schematic of the study design for the development of the prognostic marker selection and validation of the 15 prognostic markers; seven well-known biomarkers and eight newly identified features. We selected prognostic markers via the computation and overlapping of seven prognosis relevance values (PRV) lists. We chose lung adenocarcinoma (ADC) patients with complete clinical data and divided them into the training (n = 256), test (n = 171), and validation (n = 85) sets. We trained deep neural net-works (DNNs) using the training set and tuned hyper-parameters using the validation set. After training the DNNs, we classified the test set and conducted the survival analysis.

### 2.1 Identification of eight prognostic markers for feature selection

We referred seven well-known markers, which are highly correlated with NSCLC according to previous studies [5, 6, 7, 8, 9, 10, 11, 21, 22] (EPCAM, HIF1A, PKM, PTK7, ALCAM, CADM1, and SLC2A1). Based on each of these seven markers, we separated the 614 ADC samples into marker- and marker+ subgroups according to expressions by StepMiner [23].

We constructed interaction networks for both marker- and marker+ subgroups, resulting in a pair of gene interaction networks for each NSCLC marker. From each pair of interaction networks, genes were ranked by PRV (See Methods). We selected the top 30 genes as candidate markers for each well-known NSCLC marker. The details of the PRV for each marker are listed in the Supplementary Material. To guarantee robustness, we overlapped the seven PRV lists and identified markers common among all seven lists. We found that genes including COPS5, CUL1, CUL3, EGFR, ELAVL1, GRB2, HSP90AA1, NRF1, PPP1CA, RNF2, RPA2, SIRT7, and SUMO1 overlapped. By filtering these genes based on significance with p-values lower than 0.01 for survival, the final eight prognostic markers, CUL1, CUL3, EGFR, ELAVL1, GRB2, NRF1, RNF2, and RPA2, were identified.

### 2.2 Integrating gene expression and clinical data using a DNN

We used a DNN to exploit the interdependencies of the 15 selected prognostic markers, which were fed into the DNN as input features. The output of the DNN was a binary outcome of the five-year survival probability for the patient after the first therapeutic treatment. The optimized structure for our DNN uses four hidden layers, each with 40 neurons, rectified linear unit (ReLU) as the activation function, and Nadam as the optimizer. To confirm the effectiveness of the DNN on survival classification, we compared the performance with other well-known classifiers, such as K-Nearest Neighbors (KNN), Random Forest, and Support Vector Machine (SVM) by using the same prognostic markers as input features [24, 25, 26]. For the KNN classifier, the Euclidean distance was used as the distance metric, and for the SVM classifier, a Gaussian radial basis function was used as the kernel function. The parameters used in KNN, RF, and SVM were all optimized based on 10-fold cross validation of the training set. We also compared different classifiers with the molecular prognostic index (MPI) [27]. Due to the imbalance of labels across the entire data set (n = 512; survivals = 355 and deaths = 157), the classifiers tended to classify the patients as alive. In this case, if a naive classifier classified all patients as alive, it still reached an accuracy of 0.6953. To conclude, AUC is a much better performance metric than accuracy (described in the Supplementary Material). We found that the performance of the DNN (AUC: 0.7926, accuracy: 0.7485) was superior to all other methods in terms of AUC. However, the accuracy of the DNN was comparable with RF (AUC: 0.7767, accuracy: 0.7544), yet higher in AUC. Furthermore, the AUC of RF was close to our DNN model, whereby we could compare RF and existing work [27] as the candidate for further investigation. Moreover, we trained our DNN using clinical patient data (age, gender, and stage). In a previous study, a clinical prognostic index (CPI) risk score was defined by using clinical data [27]. We again noted that the DNN (AUC: 0.7388, accuracy: 0.6608) achieved a significantly higher AUC than CPI (AUC: 0.6460, accuracy: 0.6257) and RF (AUC: 0.6361, accuracy: 0.6784).

Although several studies have combined microarray and clinical data for making predictions [28], it was difficult to integrate two heterogeneous data sources. The DNN has the flexibility to integrate heterogeneous data sources by merging the hidden layers of the neural networks. The application of a DNN for integrating different types of data sources has been successful in handling audio and video data sources [29]; however, it is relatively new to integrate gene expression and clinical data sources. Therefore, we proposed using a DNN for data integration (Figure. 2a). The weights of the integrative DNNs were trained again by feeding in the microarray and clinical data simultaneously. The weights trained for the individual DNN networks were used as initial weights for the pre-training of the integrative network.

**Figure 2:**
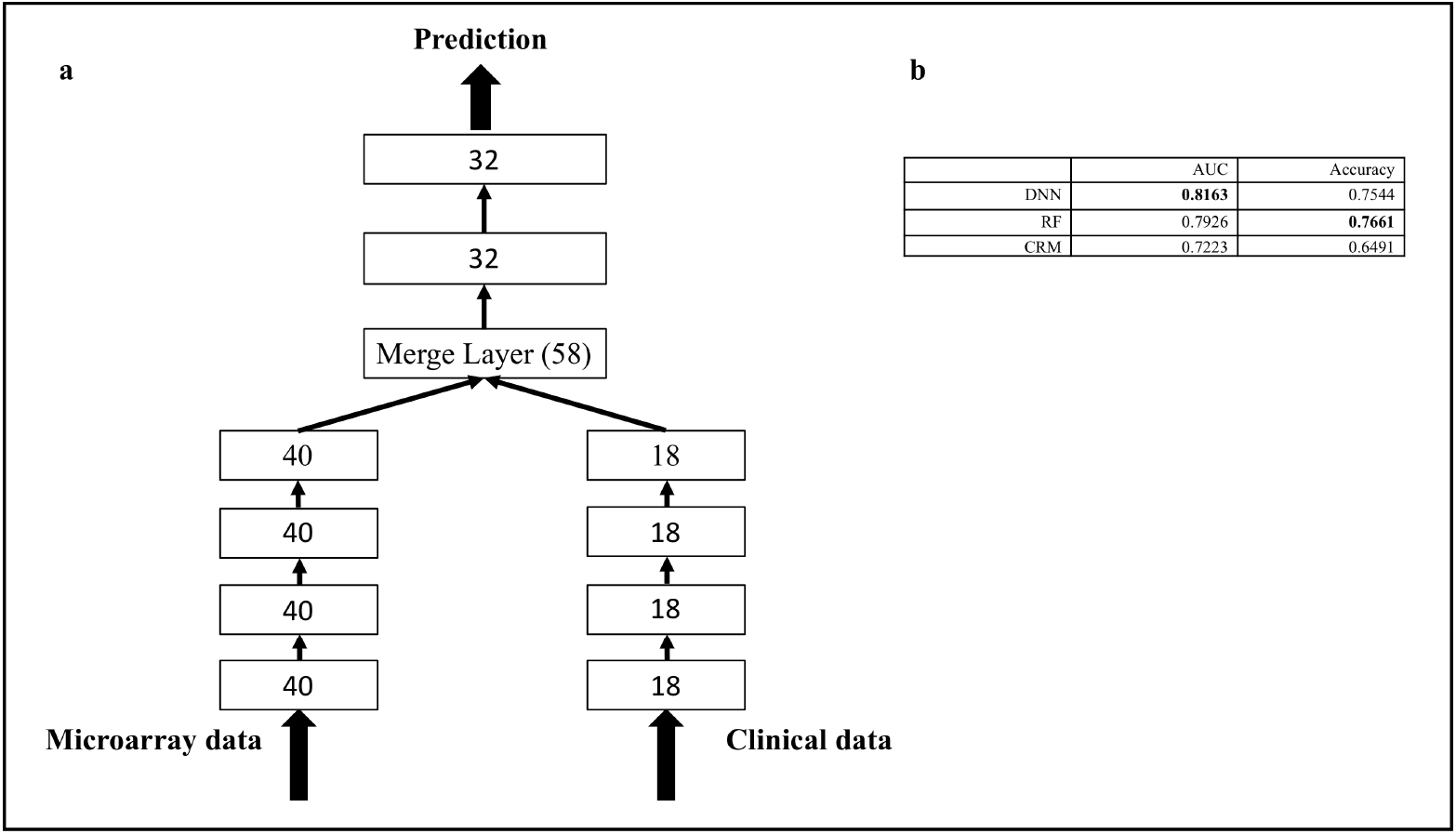
The DNN structure of combined data and comparison with other methods. (a) The left branch network deals with the microarray data source and the right branch network processes the clinical data source. Both subnetworks merge together and form an integrative network. We merged the 4th hidden layer (with 40 neurons) of the microarray DNN data and the 4th hidden layer (with 18 neurons) of the clinical DNN. The merged layer contains 58 neurons and were concatenated with two hidden layers with 32 neurons for the final prediction. (b) Comparison of the DNN with other methods for combined data.

Gentles et al. also combined gene expression data (MPI) and clinical data (CPI) to define a composite risk model (CRM). The threshold was again chosen from the median of training sets for the CRM. We further compared the performances of our proposed integrative DNN with RF and CRM, as shown in Figure. 2b.

The AUC performance of the integrative DNN (AUC: 0.8163, accuracy: 0.7544) was better than that of the RF (AUC: 0.7926, accuracy: 0.7661). It is important to note that after we included the clinical data, the improved AUC performance of the DNN (0.7926 to 0.8163, improved by 3%) was higher than that of the RF (0.7767 to 0.7926, improved by 2%). Our proposed integrative DNN is more capable of integrating heterogeneous data sources, and this was reflected by the AUC performance. Both machine learning-based algorithms significantly outperformed the CRM method (AUC: 0.7223, accuracy: 0.6491).

### 2.3 From AUC to reclassification

To obtain an overall picture of the performance comparison, we also considered precision, recall, and F1-score. The F1-score is also called the F1 measure and it considers both the precision and the recall by computing the harmonic mean of precision and recall [30]. In addition, we used the Youden index [31] as the new cut-off point for reclassification from the training set. Such reclassification was conducted for both our proposed DNN with only microarray data and the integrative DNN with both microarray and clinical data. We calculated the cut-off points (Youden indices) as 0.4396 and 0.4008 for the two DNNs, respectively. Similarly, we can also calculate Youden indices as 0.32 and 0.34 for the microarray RF and integrative RF, respectively. The cut-off points were then used as new thresholds for our DNNs and RFs. Both Youden indices are smaller than 0.5, indicating that the number of patient classified deaths increased.

To confirm the effectiveness of reclassification, we evaluated their performances in terms of accuracy, precision, recall, and F1-score for the microarray DNN and RF and the integrative DNN and RF (Figure. 3a). Note that the imbalanced structure of the data makes the recall and F1-score of the original classifier low. When Youden indices were used for reclassification, the evaluation criteria (recall and F1-score) improved for both microarray and integrative DNNs and RFs. For DNNs, we observed a significant increase in recall (from 0.3269 to 0.5961 for microarray DNN; 0.3462 to 0.7885 for integrative DNN) and F1-score (from 0.4416 to 0.5740 for microarray DNN; from 0.4615 to 0.6406 for integrative DNN). Similarly, we also observed a significant increase in recall (from 0.4230 to 0.7307 for microarray RF; 0.4423 to 0.7115 for integrative RF) and F1-score (from 0.5116 to 0.5984 for microarray RF; from 0.5349 to 0.5968 for integrative RF). For RF, we observed a minor decrease in accuracy (from 0.7485 to 0.7310 for microarray DNN; from 0.7544 to 0.7310 for integrative DNN). Furthermore, there was a decrease in precision (from 0.68 to 0.5536 for microarray DNN; from 0.6923 to 0.5394 for integrative DNN), but a significant increase in recall compensates, resulting in a significant overall improvement in the F1-score. We also observed decreases in both accuracy (from 0.7544 to 0.7017 for microarray RF; from 0.7661 to 0.7076 for integrative RF) and precision (from 0.6470 to 0.5067 for microarray RF; from 0.6765 to 0.5139 for integrative RF) for RFs. Overall, the integrative DNN achieved a higher F1-score than the integrative RF, and therefore is a more desired classifier after reclassification.

**Figure 3:**
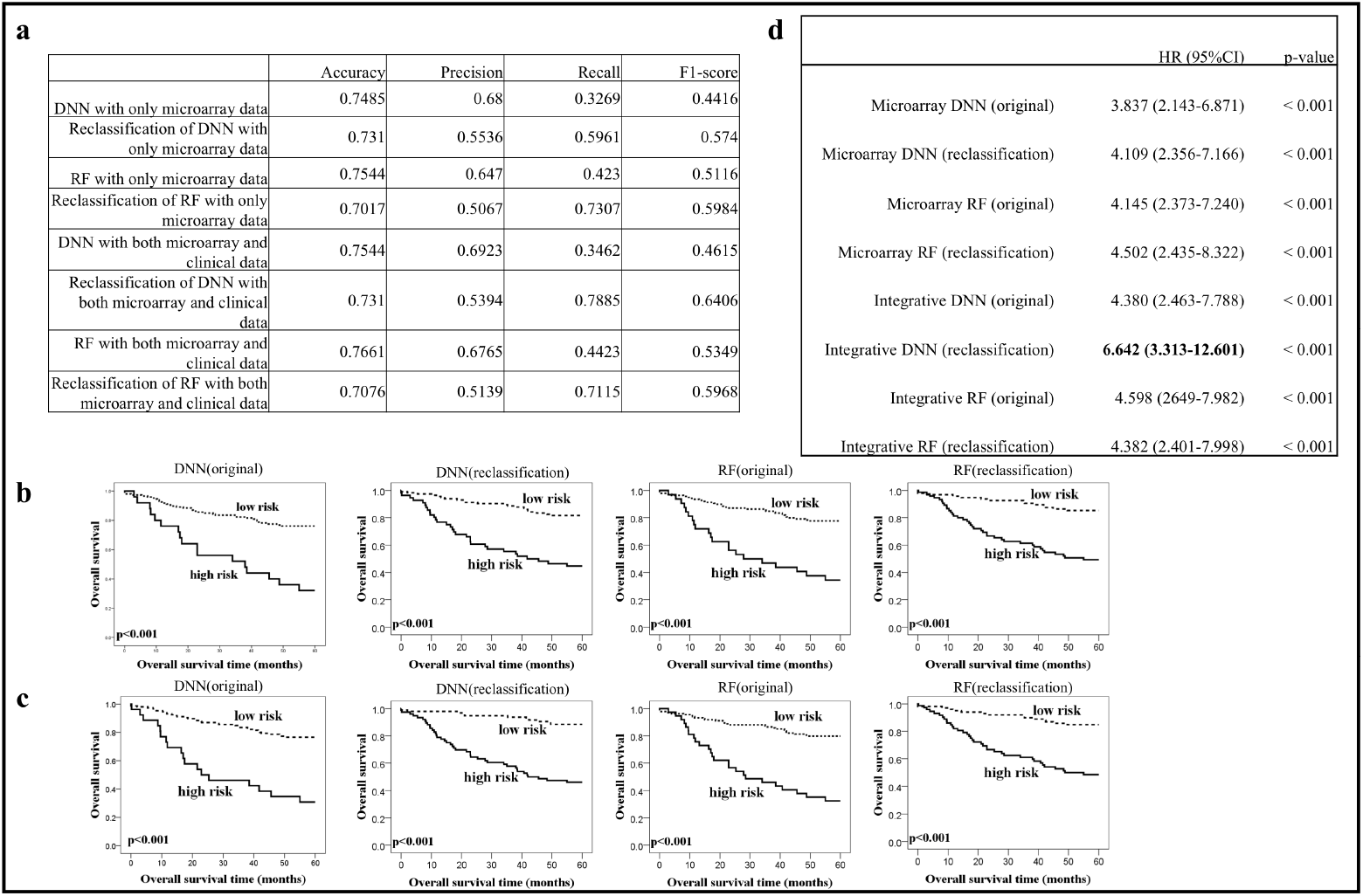
DNN performance evaluation. (a) The performance of the DNN with/without reclassification with only microarray data or both microarray and clinical data. (b) KM analysis of overall survival in the cohort microarray test set with stratification of risk groups based on the DNN and RF. Microarray data were used in our classifiers and the cut-off threshold was set at either 0.5 (original) or using the Youden index (reclassification). (c) Both microarray and clinical data were applied to classifiers and the cut-off threshold was set at either 0.5 (original) or using the Youden index (reclassification). (d) Univariate analysis of each classifier.

### 2.4 Survival analysis

To further validate the effectiveness of our proposed approach, we conducted survival analysis for ADC patients based on our prognostic markers and deep learning models. We divided the ADC patients into high risk (which was predicted as dead) and low risk (which was predicted as survived) by our proposed microarray and integrative DNNs, respectively. KM analysis (see Methods) and proportional-hazards model were used to evaluate the results for both DNN models with/without reclassification (Figure. 3). We observed that reclassification indeed separates the two risk groups further apart based on KM analysis for both our DNN models (Figure. 3b for microarray DNN; Figure. 3c for integrative DNN). An improvement can also be observed in the proportional-hazards model. The microarray DNN (original) separates patients into high and low risk groups (HR: 3.837, 95% CI: 2.143–6.871; p-value < 0.001). After reclassification, we observed a more eminent separation between the two risk groups (HR: 4.109, 95% CI: 2.356–7.166: p-value < 0.001). For the integrative DNN, the separation between the two groups becomes even more significant with reclassification (HR: 6.642, 95% CI: 3.313–12.601, p-value < 0.001). We observed similar results for RFs; however, the separation of the low and high risk groups was greater for DNNs than RFs. We can also observe that integrative DNN achieves the highest hazard ratio, indicating that bimodal DNN is capable of extracting useful information from heterogeneous data types (Figure. 3d).

### 2.5 Independent validation test

To further validate the robustness of our proposed DNN models, we evaluated their performances on an independent dataset (E-MTAB-923). The data samples for E-MTAB-923 (n = 90) are more balanced (51 survivals, 39 death) than the original cohort set. We again compared our proposed DNN models with RF in terms of AUC and accuracy (Figure. 4a). Interestingly, the integrative DNN outperformed RF not only in AUC, but also in accuracy. This suggests that our proposed integrative DNN model generalizes better to the independent validation set. For reclassification, we used the Youden indices in the previous section for microarray and integrative DNNs and RFs. Since the data samples of this validation set are more balanced than the cohort set, the results of classification are no longer extremely biased towards the low risk group. We further compared the accuracy, precision, recall, and F1-score of the microarray DNN and RF and the integrative DNN and RF (Figure. 4e). Due to different label distributions between training set and independent validation set, we observed that reclassification lowered the accuracies of both DNNs (from 0.6556 to 0.5889 for microarray DNN; from 0.6899 to 0.6111 for integrative DNN). However, reclassification improved the recall and F1-score in both DNNs. We had similar results for RFs. For RFs, we also observed an increasing trend in recall (from 0.2307 to 0.5897 for microarray RF; 0.1795 to 0.6923 for integrative RF) and F1-score (from 0.3529 to 0.5679 for microarray RF; from 0.2917 to 0.6353 for integrative RF). Although reclassification decreased accuracy, it also improved F1-score. The integrative DNN model has the best F1-score and AUC in the independent validation set.

**Figure 4:**
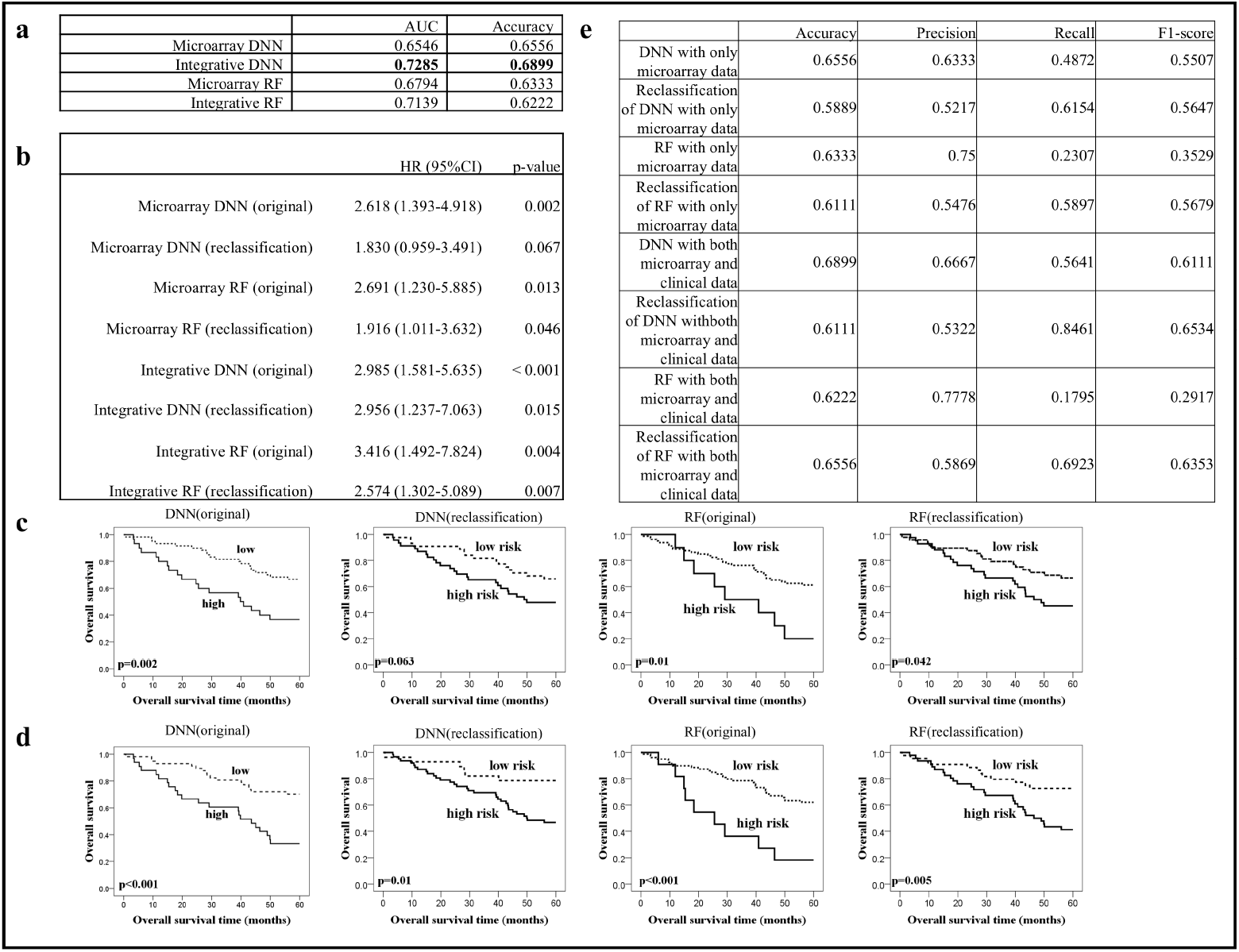
Survival analysis. (a) The performance of the DNN and RF on the independent validation set. (b) Univariate analysis of each classifier. (c) KM analysis of overall survival in the independent validation set with stratification of risk groups based on the DNN and RF. Microarray data were used in our classifiers and the cut-off threshold was set at either 0.5 (original) or using the Youden index (reclassification). (d) Both microarray and clinical data were applied to classifiers and the cut-off threshold was set at either 0.5 (original) or using the Youden index (reclassification). (e) The performance of the DNN with/without reclassification with only microarray data or both microarray and clinical data.

We again divided the independent validation set into two risk groups and used the KM analysis and proportional-hazards model for survival analysis (Figure. 4). Although the stratifications did not benefit from adding clinical data, improvement was observed for the proportional-hazards model. The microarray DNN (original) separated patients into high and low risk groups (HR: 2.618, 95% CI: 1.393–4.918; p-value = 0.002). After reclassification, we observed that the separation between the two risk groups was closer (HR: 1.830, 95% CI: 0.959–3.491; p-value = 0.067). For the integrative DNN, the separation between the two groups becomes even more eminent (HR: 2.985, 95% CI: 1.581–5.635; p-value < 0.001). On the other hand, we observed a moderate separation for the microarray RF (original) (HR: 2.691, 95% CI: 1.230–5.885; p-value = 0.013). After including both microarray and clinical data, we observed a more significant separation between the two risk groups (HR: 3.416, 95% CI: 1.492–7.824; p-value = 0.004). We expected the results from survival analysis to be worse since the reclassification reduced accuracy. Recall of both DNNs was improved, although the separation between the two groups was still not eminent after reclassification. However, this was a trade-off between F1-score and accuracy.

## 3 Discussion

Traditionally, feature selection methods can be categorized into three different types: the filter, wrapper, and embedded methods [32]. The strategy for the filter method was based on ranking the performance or mutual information of each feature, which was widely used in many domains because of their simplicity. The wrapper method worked as a black box by selecting features based on the performance of the classifier, which often showed great computational complexity and only worked well in limited classifiers. The embedded method was similar to the wrapper method; however, it focused on reducing the amount of computational time required. All of the methods above aimed to select the top-ranked features purely based on statistical or predictive performance and disregarded the biological meaning of the gene features. Therefore, the selected gene features often lack biological insights and cannot be applied to further experimental validation.

To address this issue, we demonstrated a systems biology approach to select biologically meaningful gene features and identify eight prognostic markers for NSCLC patients. These prognostic markers (CUL1, CUL3, EGFR, ELAVL1, GRB2, NRF1, RNF2, and RPA2) were identified by overlapping seven computed PRV lists. Among the eight prognostic markers, most genes have been reported and directly relate to NSCLC in previous studies. ELAVL1 is a well-known RNA-binding protein associated with multi-carcinogenesis, such as large cell lymphoma and glioma, by modulating RNA stability [33]. Overexpression of nuclear ELAVL1 in NSCLC patients was correlated with lymph node metastasis and may serve as a potential diagnosis biomarker [34]. In addition, while nuclear ELAVL1 was important in cancer progression, cytoplasmic ELAVL1 was also identified as an independent prognostic factor for survival in NSCLC [35]. EGFR is a well-known transmembrane protein involved in controlling cell survival and tumorigenesis in many malignancies, including NSCLC [36]. Moreover, mutated and overexpressed EGFR has been reported in a myriad of NSCLC case studies [37]. Interestingly, while EGFR was recognized as a potential therapeutic target of NSCLC, its binding adaptor, growth factor receptor bound protein 2 (GRB2), was shown to be a stabilized EGFR and co-activated downstream to the MAPK/ERK pathway [37]. Among the Cullion family, Cullin 1 (CUL1) is one of the scaffold proteins in E3 ubiquitin ligase involved in cancer progression. High expression was correlated to patient survival rate, which was identified as an independent prognosis factor in NSCLC [38]. There are eight members in the Cullin family. In addition to CUL1, CUL3 is known as a scaffold protein in the ubiquitin-proteasome system and contributes to cellular regulation, such as cell cycles, protein trafficking, and stress response, which are common tumorigenesis phenomenon when mutated. Furthermore, one substrate adaptor of CUL3, kelch-like ECH-associated protein (Keap), was first identified as an inhibitor of transcriptional factor Nf-E2-related factor 2 (Nrf2) and played important roles in anti-oxidation stress and cell defense during cancer suppression [39]. Although there is only limited evidence addressing the function of Nrf1 and Nrf2 in prostate cancer [40], the genetic and functional conservation between them identifies their active roles in lung cancer progression. Ring finger protein 2 (RNF2), a member of the group II polycomb group (PcG) protein, was highly expressed in many types of human malignancies [41]. Despite the role RNF2 plays in biological processes in cancers via its diverse mechanisms, it highlights the potential oncogenic activity of RNF2 on NSCLC. Replication protein A 2 (RPA2), a single-strand DNA binding protein, processes DNA metabolism in response to DNA damage-associated replication arrest [42]. It has also been considered a potential therapeutic target and prognosis indicator for colon cancer, as shown by differences in its immunohistochemical expression between cancer patients and controls [43]. To summarize, each marker we identified showed great potential for being a prognosis biomarker based on its biological background.

We further analyzed eight prognostic markers using GSEA (Gene-Set Enrichment Analysis) [44]. Using GO biological process enrichment analysis, we found that CUL1, CUL3, RNF2, and RPA2 overlap in their mitotic cell cycles, which has important biological implications in tumor development. We also analyzed these prognostic markers by investigating their interdependencies using STRING (https://string-db.org/) [45] (Figure. 5). The interaction between GRB2 and EGFR were experimentally verified [46]; both were characterized in the KEGG “Non-small cell lung cancer pathway” (pathway #5223). In addition, the interactions among GRB2, EGFR, HIF1A, and SLC2A1 were also highly correlated with cancer in the KEGG “Pathways in cancer” (pathway #5200).

**Figure 5:**
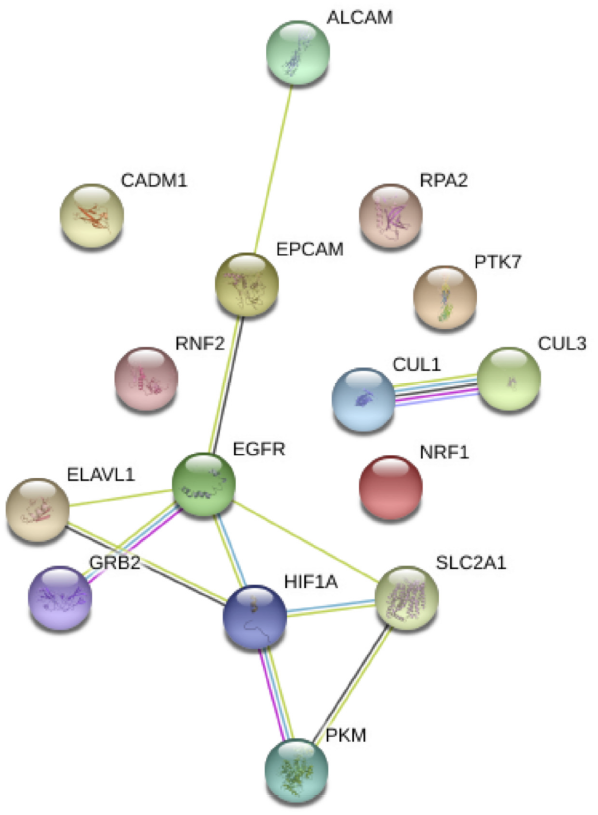
The interaction network of prognostic markers.

One reason our proposed PRV feature selection method performs well in not only the test set, but also the independent validation set, is due to the robustness emphasized in our systems biology approach. Several steps were taken to ensure the robustness of our feature selection method. Firstly, we used gene expression data from six different datasets and selected only genes with the same probe number. Secondly, our PRV feature selection method was applied to each of the seven well-known NSCLC markers in our data cohort. We then constructed seven different paired interaction networks to obtain seven PRV lists. The eight prognostic marker genes were selected via overlapping these seven PRV lists. In other words, we only selected the prognostic markers that showed in all seven PRV lists corresponding to the seven well-known NSCLC markers. This selection process guarantees robustness in our method based on the superior predictive performance of our proposed integrative DNN. In the future, we can not only choose well-known markers from the literature, but use other clinical outcomes to group samples for selecting features.

In most of the existing work, only gene expressions were used as features for training classifiers, with the aim of predicting various disease outcomes [12, 14]. In recent years, some researchers have begun to consider combining gene expression and clinical data to make such predictions [27, 28]. In this study, we applied the concept of “bimodal learning” to construct an integrative DNN where two heterogeneous modalities (gene expression and clinical data) were integrated for predicting ADC patient prognosis. By using two modalities, the integrative DNN approach is capable of providing the missing information left by the other observed modality. Compared with our microarray DNN, we observed an increase in AUC and accuracy from the integrative DNN. We also demonstrated improved prognosis performance for survival analysis. This highlights the benefit of integrating microarray and clinical data via our integrative DNN approach. A good bimodal learning model possesses certain properties. The joint representation should be similar in its feature space, implying that the two heterogeneous data sources correspond to similar concepts. We use two DNNs to extract features from each data source and jointly train them for the complete bimodal learning model in the combined layers. From this, the combined model better integrates two data modalities into a joint representation. For future work, we could possibly use more than two types of data sources to construct a multimodal learning model for more accurate classification.

## 4 Methods

### 4.1 Microarray data preprocessing

The open access microarray data were downloaded from the National Center for Biotechnology Information (NCBI) Gene Expres-sion Omnibus (GEO) database (http://www.ncbi.nlm.nih.gov/geo) with accession numbers GSE19188, GSE29013, GSE30219, GSE31210, GSE37745, GSE50081 using the same platform: the human Affymetrix HG-U133 Plus 2.0 platform (GPL570). Six independent GEO datasets were merged into one cohort set with 614 samples for microarray analysis. We separated 256 samples as the training set, 85 samples as the validation set, and 171 samples as the test set, all of which have complete clinical data. E-MTAB-923 was taken as an independent validation set with the same platform. Table 1 provides detailed information on these datasets. The median overall survival time is 56.25 months. All the datasets were preprocessed by the robust multi-array average (RMA) algorithm and gene expression values were log2 transformed.

**Table 1:**
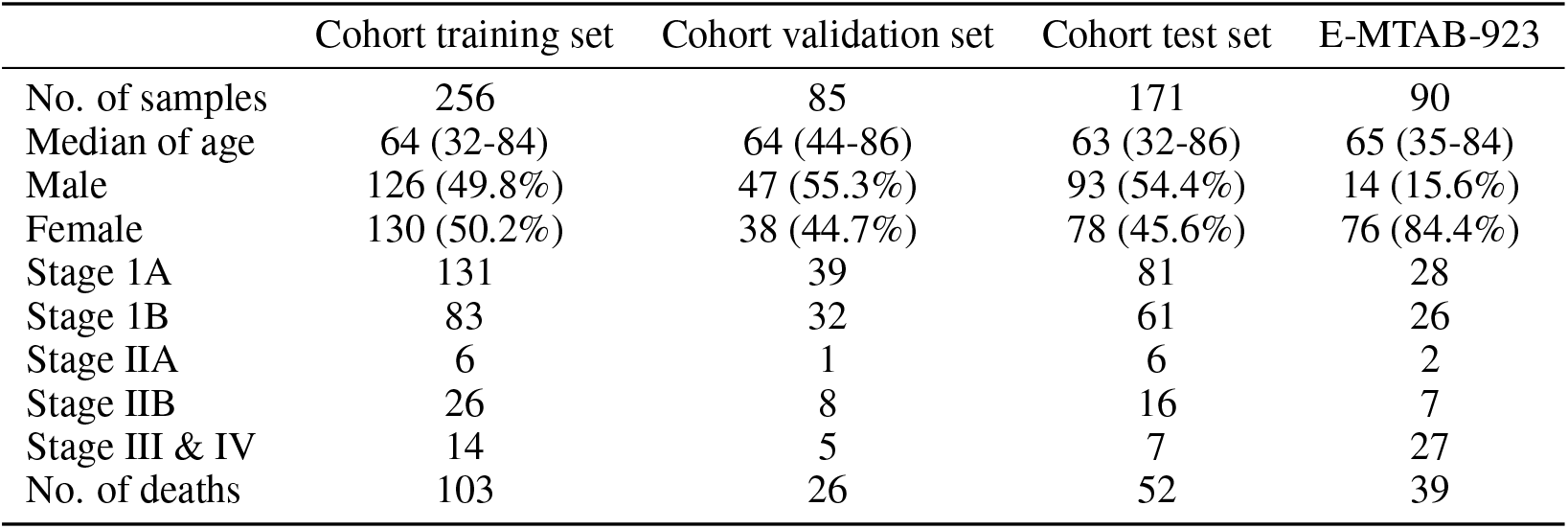
Clinical characteristics of patients.

|  | Cohort training set | Cohort validation set | Cohort test set | E-MTAB-923 |
| --- | --- | --- | --- | --- |
| No. of samples | 256 | 85 | 171 | 90 |
| Median of age | 64 (32-84) | 64 (44-86) | 63 (32-86) | 65 (35-84) |
| Male | 126 (49.8%) | 47 (55.3%) | 93 (54.4%) | 14 (15.6%) |
| Female | 130 (50.2%) | 38 (44.7%) | 78 (45.6%) | 76 (84.4%) |
| Stage 1A | 131 | 39 | 81 | 28 |
| Stage 1B | 83 | 32 | 61 | 26 |
| Stage IIA | 6 | 1 | 6 | 2 |
| Stage IIB | 26 | 8 | 16 | 7 |
| Stage III & IV | 14 | 5 | 7 | 27 |
| No. of deaths | 103 | 26 | 52 | 39 |

### 4.2 Prognosis relevance value

Based on the gene expression for each of the well-known markers, we divided patients into two different subgroups (marker-and marker+) via the StepMiner algorithm [23]. Crucial genes were identified by a systematic comparison between subgroups. For both marker- and marker+ groups, we constructed the corresponding gene interaction networks (interaction networks) [47]. According to the constructed interaction networks, we defined prognosis relevance values (PRV) to measure the difference between marker+ and marker-interaction networks for each gene (details in the Supplementary Material). Genes with a larger PRV show a significant difference among interactions or connections compared with other genes, which are potential prognostic markers.

### 4.3 Bimodal DNN

A DNN is composed of one input and one output layer, with many hidden layers in between representing multiple levels of abstraction. Each hidden layer is composed of many neurons. Deep learning has been successfully applied to supervised learning for combining different modalities. For our dataset, we not only used microarray data, but also clinical data. Here, we combined two DNNs (one for the microarray data input and the other for the clinical data input) by merging their output layers and further concatenating several hidden layers before reaching the final decision. The integrative network can be expected to benefit from combining the two separate data sources.

### 4.4 From AUC to reclassification

The receiver operating characteristic [48] curve is a graphical plot that illustrates the diagnostic ability of a binary classifier system created by plotting the true positive rate against the false positive rate at various threshold settings. The area under the ROC curve has been used for diagnostic testing in radiology [48]. Under some conditions, we can achieve a better performance by adjusting the cut-off points. In this study, the cut-off points were determined by the Youden index [31], which is a frequently used summary measure of the ROC curve. In previous classification tasks, we classified a sample by comparing the probabilities of patient survival outcomes. To further improve predictive performance, new cut-off points to be determined are used for reclassification.

### 4.5 Survival analysis

In our study, overall survival time was calculated from the date of surgery to the date of death. We predicted the survival rate of patients after the 5-year prognosis. Therefore, all patients were treated as alive samples when survival time was greater than five years. Survival curves were demonstrated based on a Kaplan-Meier estimation for five years and compared using a log-rank test [49] (KM analysis). We applied the cox proportional-hazards model to analyze the relationship between the prognostic genes for survival [50]. The hazard ratios (HR) and confidence intervals (CI) were reported.

## Acknowledgements

We would like to thank Uni-edit (www.uni-edit.net) for editing and proofreading this manuscript.

## Author contributions

YHL, WNC, and CL conceived of the conception and design of this research. YHL, WNC, CL, and YT developed methodologies. WNC and CL are responsible for acquisition of data (provided animals, acquired and managed patients, provided facilities, etc.). YHL, WNC, CL, and TCH performed analyses and interpreted data (e.g., statistical analysis, biostatistics, computational analysis). YHL, WNC, CL, TCH, YT, and SW wrote, reviewed, and/or revised the manuscript. YHL, WNC, CL, TCH, YT, and SW provided administrative, technical, or material support (i.e., reporting or organizing data, constructing database). CL, YT, and SW supervised this study. All authors read and approved the final manuscript.

## Competing interests

The authors declare no competing interests.

## Data availability

All data generated or analyzed during this study are included in this published article and its supplementary information files.

