## Supplementary for "Predicting the Prognosis of Non-Small Cell Lung Cancer by Integrating Microarray and Clinical Data with Deep Learning"

### Calculation of prognostic protein relevance value

Based on the gene expression of each of the seven well-known markers, we could divide patients into two different subgroups, marker- and marker+, via the StepMiner algorithm to identify crucial genes using a systematic comparison between these subgroups. The goal of the StepMiner algorithm is to find a one-step function that fits the ordered samples. The StepMiner algorithm defined a “step” for a marker according to the largest jump of expression values from low-to-high as a threshold. To suppress noise, an intermediate region was defined as 0.5 below and 0.5 above the threshold. We then obtained two subgroups for each well-known marker, i.e., the marker- and marker+ groups, excluding the noise zone. For both the marker- and marker+ groups, we constructed the corresponding interaction networks. To construct the interaction network, the following dynamic model was defined for the i-th protein:

$$x_i[n] = \sum_{j=1}^{Ni} a_{ij} x_j[n] + \omega_i[n] \tag{1}$$

where $x_i[n]$ denotes the expression levels for the i-th protein at sample n; $a_{ij}$ denotes the interactions ability between the i-th protein and its j-th interacting protein; $\mathrm{N_i}$ is the total number of PPIs with the i-th proteins; and $\omega_i[n]$ represents stochastic noise. In order to prune the false positive interactions, we applied the Akaike information criterion (AIC) model order selection and t-test to reveal the number of valid associations and determine the model parameters.

The dynamic model for the interaction network in (1) can be rewritten as:

$$X(n) = AX(n) + \omega(n) \tag{2}$$

where

$$X(n)=\begin{bmatrix} x_1[n] \\ x_2[n] \\ \vdots \\ x_M[n] \end{bmatrix},\ A=\begin{bmatrix} a_{11} & \cdots & a_{1M} \\ \vdots & a_{ij} & \vdots \\ a_{M1} & \cdots & a_{MM} \end{bmatrix},\ W(n)=\begin{bmatrix} \omega_1[n] \\ \omega_2[n] \\ \vdots \\ \omega_M[n] \end{bmatrix}$$

where $M$ is the number of interacting proteins in the refined interaction network; matrix $A$ denotes the interactions ability matrix; and $W(n)$ is the noise matrix.

If $a_{ij}$ is equal to zero, then there is no interaction between protein i and protein j or it is identified as a false positive and pruned in the refined interaction network. For convenience, $A_k^+$ and $A_k^-$ are denoted as the interaction ability matrices for marker+ and marker- interaction networks, respectively:

$$A_k^+=\begin{bmatrix} a_{11,k}^+ & \cdots & a_{1M,k}^+ \\ \vdots & \ddots & \vdots \\ a_{M1,k}^+ & \cdots & a_{MM,k}^+ \end{bmatrix},\ A_k^-=\begin{bmatrix} a_{11,k}^- & \cdots & a_{1M,k}^- \\ \vdots & \ddots & \vdots \\ a_{M1,k}^- & \cdots & a_{MM,k}^- \end{bmatrix}$$

where k denotes the index for the considered well-known marker.

According to the constructed interaction networks, we define a matrix $D_k$ to measure the difference between marker+ and marker- interaction networks for the k-th well-known marker:

$$D_k = A_k^+ - A_k^- = \begin{bmatrix} d_{11,k} & \cdots & d_{1M,k} \\ \vdots & \ddots & \vdots \\ d_{M1,k} & \cdots & d_{MM,k} \end{bmatrix} \tag{3}$$

$$= \begin{bmatrix} a_{11,k}^+ - a_{11,k}^- & \cdots & a_{1M,k}^+ - a_{1M,k}^- \\ \vdots & \ddots & \vdots \\ a_{M1,k}^+ - a_{M1,k}^- & \cdots & a_{MM,k}^+ - a_{MM,k}^- \end{bmatrix} \tag{4}$$

where $d_{ij,k}$ denotes the difference in interaction abilities between the i-th and j-th protein for the k-th well-known marker. To identify candidate prognostic markers from $D_k$, the PRV is defined as follows:

$$PRV_{i,k} = \sum_{j=1}^{M} \left| d_{ij,k} \right| \tag{5}$$

where $PRV_{i,k}$ in (5) represents the PPI difference for the i-th protein and the k-th well-known marker.

### PRV lists

#### Table S1: Top 30 PRV proteins for seven well-known NSCLC markers

| EPCAM | HIF1A | PKM | PTK7 | ALCAM | CADM1 | SLC2A1 |
|---:|---:|---:|---:|---:|---:|---:|
| UBC | UBC | **ELAVL1** | UBC | UBC | UBC | UBC |
| **NRF1** | **NRF1** | **NRF1** | **NRF1** | **ELAVL1** | **NRF1** | **NRF1** |
| **ELAVL1** | **ELAVL1** | APP | **ELAVL1** | **NRF1** | **ELAVL1** | **ELAVL1** |
| APP | **SUMO1** | **SUMO1** | APP | APP | APP | HSPA4 |
| **SUMO1** | **RNF2** | **RNF2** | **SUMO1** | **SUMO1** | **SUMO1** | **SUMO1** |
| **CUL3** | **CUL3** | **CUL3** | **CUL3** | **CUL3** | **CUL3** | HEC W2 |
| **RNF2** | **SIRT7** | NEDD8 | **SIRT7** | HNRNPU | **RNF2** | **CUL3** |
| **COPS5** | NEDD8 | **COPS5** | HSPA4 | HECW2 | **COPS5** | NEDD8 |
| **EGFR** | **EGFR** | **RPA2** | HECW2 | **RNF2** | YWHAQ | **RNF2** |
| **HSP90AA1** | **GRB2** | **SIRT7** | **RPA2** | **SIRT7** | **GRB2** | ILF3 |
| PARK2 | **RPA2** | **GRB2** | **RNF2** | **COPS5** | **RPA2** | **COPS5** |
| **GRB2** | **COPS5** | YWHAQ | NEDD8 | NEDD8 | **SIRT7** | HNRNPU |
| HECW2 | **HSP90AA1** | **HSP90AA1** | **COPS5** | ILF3 | **EGFR** | SIRT1 |
| **SIRT7** | YWHAB | **EGFR** | CALM1 | **EGFR** | **HSP90AA1** | **HSP90AA1** |
| DHX9 | **PPP1CA** | PARK2 | **GRB2** | **GRB2** | PARK2 | **SIRT7** |
| SRRM2 | ICT1 | **CUL1** | **HSP90AA1** | **RPA2** | YWHAB | **GRB2** |
| YWHAQ | **CUL1** | YWHAB | WWOX | **HSP90AA1** | IGSF8 | YWHAB |
| CDKN1A | RPA1 | CUL5 | HNRNPU | **PPP1CA** | **PPP1CA** | **PPP1CA** |
| YWHAB | CAND1 | HDAC1 | CDKN1A | YWHAQ | **CUL1** | SMAD3 |
| **PPP1CA** | HDAC1 | FBXO6 | PARK2 | SIRT1 | HECW2 | **RPA2** |
| VCP | VCP | HDAC2 | YWHAQ | YWHAB | DHX9 | **EGFR** |
| **CUL1** | CDK2 | MDFI | **CUL1** | **CUL1** | CALM1 | YWHAQ |
| CALM1 | KDM1A | BMI1 | EIF4A3 | SMAD3 | FBXO6 | HSPA8 |
| H2AFX | FN1 | KDM1A | RB1 | PARK2 | CDKN1A | CREBBP |
| FBXO6 | CUL5 | **PPP1CA** | FBXO6 | FBXO6 | HDAC6 | **CUL1** |
| HDAC1 | CUL2 | IKBKG | HDAC6 | CAND1 | HDAC1 | PARK2 |
| CREBBP | COPS6 | HDAC3 | **PPP1CA** | HDAC1 | CDK2 | HDAC6 |
| HDGF | PAXIP1 | HDAC6 | **EGFR** | IKBKG | VCP | FBXO6 |
| **RPA2** | ILF3 | ICT1 | CREBBP | ICT1 | ILF3 | HDAC1 |
| HNRNPU | RPA3 | CDK2 | SMAD2 | VCP | HDGF | HDAC2 |

### Survival analysis for the eight identified prognostic gene markers

We further conducted survival analysis for our eight identified prognostic markers (CUL1, CUL3,

EGFR, ELAVL1, GRB2, NRF1, RNF2, and RPA2). Based on the expression level of each prognostic feature gene, patients were stratified into two groups, marker- and marker+, by StepMiner. The survival analysis was conducted using Kaplan-Meier methods and a univariate proportional-hazards model, as illustrated in Table S2 and Figure S1. This indicates that all of the eight prognostic feature genes were already feasible for prognosis to a certain degree. Cancer, however, is a complicated disease. It is unlikely that a single prognostic marker can accurately predict the clinical outcome of a cancer patient. We therefore exploited the interdependencies between the eight prognostic markers combined with the original seven well-known NSCLC markers via a DNN. Hopefully, we can provide a more accurate and robust approach for predicting the prognosis of NSCLC patients.

**Table S2: Univariate analysis of each prognostic marker**

| | HR (95% CI) | p-value |
|---|---|---|
| CUL1 | 1.166 (0.854–1.591) | 0.334 |
| CUL3 | 0.761 (0.573–1.012) | 0.061 |
| EGFR | 0.319 (0.227–0.448) | < 0.001 |
| ELAVL1 | 0.330 (0.232–0.469) | < 0.001 |
| GRB2 | 1.386 (0.683–2.814) | 0.366 |
| NRF1 | 0.400 (0.298–0.537) | < 0.001 |
| RNF2 | 0.671 (0.506–0.890) | 0.006 |
| RPA2 | 1.040 (0.550–1.965) | 0.904 |

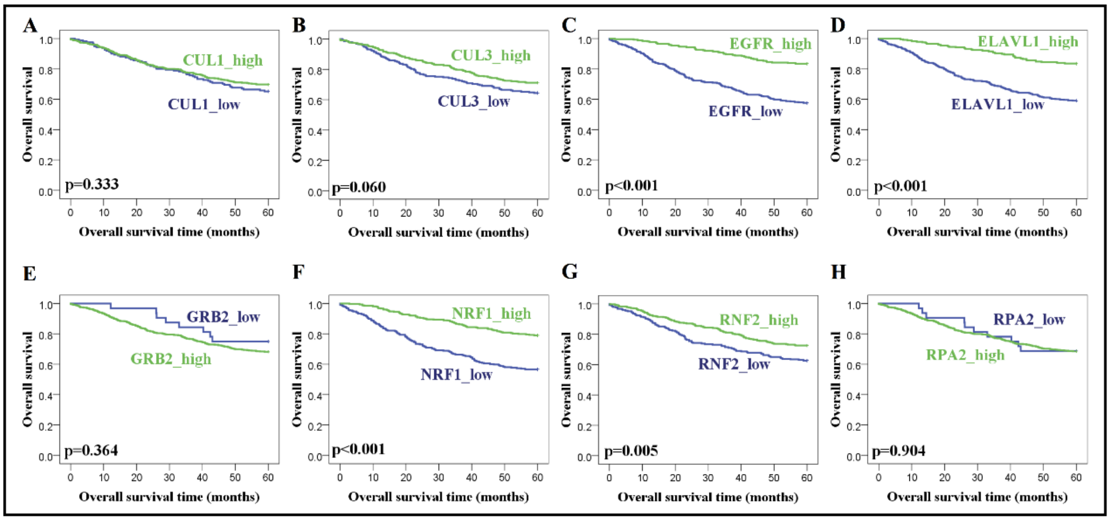

**Figure S1**. **KM curves of the eight prognostic markers in the cohort dataset.** To study the relationship between prognostic markers expression and survival, we used the StepMiner algorithm to stratify the NCBI-GEO datasets into two groups. (A–H) NSCLC patients (n = 614) were divided into two groups, which were analyzed for overall survival.

### Using DNN to exploit the interdependencies of the selected prognostic gene markers

In this section, we used a DNN to exploit the interdependencies of the 15 selected prognostic markers (seven well-known and eight newly identified markers). Specifically, these 15 markers of a patient were fed into the DNN as input features. The output of the DNN was a binary outcome of survival probability of the patient five years after the first treatment. We used 512 ADC patient samples to train the DNN, of which half were randomly assigned as a training set (n = 256), a third of the samples as a test set (n = 171), and the remaining sixth as a validation set (n = 85). The DNN was determined based on the training set. After training, we optimized classification performance by adjusting the DNN hyper-parameters (numbers of hidden layers and neurons in each hidden layer) based on the validation set. We evaluated the performance of the DNN for different hidden layers (e.g. 1, 2, 3, 4, and 5) with a different number of neurons (e.g. 8, 16, 24, 32, 40, and 48) via the validation set.

Due to the small sample size of the validation set, we obtained almost the same accuracy in several different structures, making it difficult to determine the best structure. Therefore, we instead evaluated DNN model performance using the loss function (cross entropy). We concluded that when the number of neurons is small, the number of hidden layers reduces the loss function. As indicated in Table S3, the DNN structure that comprises four hidden layers, with 40 neurons per layer, achieves minimum loss. This DNN structure was used in the following experiments.

**Table S3: Loss function of DNN with different numbers of layers and neurons**

| neurons / layers | 8 | 16 | 24 | 32 | 40 | 48 |
|---:|---:|---:|---:|---:|---:|---:|
| 1 | 0.6068 | 0.6115 | 0.5605 | 0.5505 | 0.5683 | 0.5544 |
| 2 | 0.5653 | 0.5791 | 0.5426 | 0.5828 | 0.5601 | 0.5503 |

| | | | | | | |
|---|---|---|---|---|---|---|
| 3 | 0.5476 | 0.5626 | 0.5463 | 0.5829 | 0.5415 | 0.5564 |
| 4 | 0.5479 | 0.5957 | 0.5735 | 0.5615 | **0.5361** | 0.5570 |
| 5 | 0.5392 | 0.5634 | 0.5558 | 0.5556 | 0.5541 | 0.5549 |

We further tested the activation functions, such as Sigmoid, hyperbolic tangent (Tanh), and rectified linear unit (Relu), with different optimizers (SGD, Adam, Adamax, and Nadam) on the survival classification task of the DNN. The results are summarized in Table S4.

**Table S4: Loss function of DNN with different activation functions and optimizers**

| | Sigmoid | Tanh | Relu |
|---|---|---|---|
| SGD | 0.6125 | 0.5778 | 0.5610 |
| Adam | 0.5476 | 0.5885 | 0.5461 |
| Nadam | 0.5463 | 0.5525 | **0.5361** |
| Adamax | 0.5476 | 0.5920 | 0.5521 |

We observed that the optimizer Nadam was uniformly better than SGD for the same DNN structure for each of the tested activation functions. When we used Nadam as the preferred optimizer, the loss function was smallest for the Relu activation functions. After extensive numerical experiments, the final model structure of our DNN contained four layers, with 40 neurons in each layer, along with the Relu activation function and Nadam optimizer.

### Comparison of DNN with other classifiers

The threshold was chosen from the median of the training sets for MPI. In Table S5, we compared the DNN with all the above methods by evaluating their performance on the test set. We demonstrated that the performance of the DNN (AUC: 0.7926, accuracy: 0.7485) was superior to all other methods in terms of AUC. Note that the DNN has comparable accuracy with RF (AUC: 0.7767, accuracy: 0.7544), but a higher AUC. We observed that the MPI has the worst performance both in AUC and accuracy. This indicates that our proposed framework that integrates systems biology and deep learning approaches to predict prognosis of NSCLC is more favorable than the MPI approach, even when the DNN is replaced by other well-known classifiers.

**Table S5: Comparison of DNN with other methods for microarray data**

| | AUC | Accuracy |
|---:|---:|---:|
| DNN | **0.7926** | 0.7485 |
| RF | 0.7767 | **0.7544** |
| KNN | 0.7705 | 0.7135 |
| SVM | 0.7275 | 0.7018 |
| MPI | 0.7209 | 0.6725 |

We also considered different gene features, e.g., seven well-known NSCLC markers, eight identified prognostic markers, and nine MPI markers for the microarray DNN (**Table S6**). We observed that irrespective of whether the classifier used seven well-known markers or eight newly prognostic markers as features, the predictions for both were poorer than when nine MPI markers were used. However, the classifier had the best prediction when we included seven well-known markers and eight newly prognostic markers as features.

**Table S6: Other gene features for microarray DNN**

| | AUC | Accuracy |
|---:|---:|---:|
| Seven well-known + eight newly | 0.7926 | 0.7485 |
| Nine MPI markers | 0.7736 | 0.7017 |
| Seven well-known markers | 0.7314 | 0.6959 |
| Eight identified markers | 0.7233 | 0.6842 |

### Training DNN using clinical data

In this section, we trained our DNN using patients' clinical data (age, gender, and stage). Hyper-parameters were tuned the same way as described in the previous section. The optimized structure for our DNN uses five hidden layers, each with 18 neurons, Relu as the activation function, and Nadam as the optimizer. Gentles et al. also defined a clinical prognostic index (CPI) risk score using patient clinical data. RF and CPI are compared in Table S7.

**Table S7: Comparison of DNN with other methods for clinical data**

| | AUC | Accuracy |
|---|---|---|
| DNN | **0.7388** | 0.6608 |
| RF | 0.6361 | **0.6784** |
| CPI | 0.6460 | 0.6257 |

We can again see that our DNN (AUC: 0.7388, accuracy: 0.6608) achieves significantly higher AUC than the other methods and a comparable accuracy with RF (AUC: 0.6361 accuracy: 0.6784). This may indicate that our DNN is more capable of capturing the complicated interdependencies between the features of the clinical data with the cancer survival outcome.

### Different data fusion methods

We also considered other data fusion methods for the DNN; Early Fusion (concatenated all features in first layer), Intermediate Fusion (bimodal learning), and Late Fusion (merging fully connected two sub-networks' output layers), as shown in Table S8. We can observe that the Intermediate Fusion method had the best performance.

**Table S8: Other data fusion methods for combined DNN**

| | AUC | Accuracy |
|---|---|---|
| Early Fusion | 0.8000 | 0.7368 |
| Intermediate Fusion | 0.8163 | 0.7544 |
| Late Fusion | 0.7958 | 0.7544 |

### Different data partition

To confirm that our data is not limited to one special partition, we tested our model on three different data partitions. We divided the dataset into three partitions, then took the first (original), second, and third partition as test sets in each round, respectively (Table S9 and Table S10). Although the second and third results were not as good as previously shown, they still showed acceptable classification capabilities.

**Table S9: Microarray DNN for 3-fold partition**

| | AUC | Accuracy |
|---:|---:|---:|
| Original | 0.7926 | 0.7485 |
| Second | 0.7593 | 0.7368 |
| Third | 0.7606 | 0.7000 |
| Mean | 0.7708 | 0.7284 |

**Table S10: Combined DNN for 3-fold partition**

| | AUC | Accuracy |
|---:|---:|---:|
| Original | 0.8163 | 0.7544 |
| Second | 0.7799 | 0.7836 |
| Third | 0.7696 | 0.7294 |
| Mean | 0.7886 | 0.7554 |

### Discussion of death probability for patients

We discussed probability regions of patients who had either survived or died. There are three regions ([Probability < Youden index], [Youden index < Probability < 0.5], and [0.5 < Probability]) in our discussion. We observed that it was difficult for the classifiers to classify patients in the [Youden index < Probability < 0.5] region (Table S11 and Table S12). The probability distribution is also plotted in Figure S2.

**Table S11: The distribution of patients in three regions for microarray DNN in the test set.**

| Probability region | Survival | Death |
|---:|---:|---:|

| | | |
|---|---|---|
| Probability<Youden index | 94 | 21 |
| Youden index<Probability<0.5 | 17 | 14 |
| 0.5<Probability | 8 | 17 |

**Table S12: The distribution of patients in three regions for integrative DNN in the test set.**

| Probability region | Survival | Death |
|---|---|---|
| Probability < Youden index | 84 | 11 |
| Youden index < Probability < 0.5 | 27 | 23 |
| 0.5 < Probability | 8 | 18 |

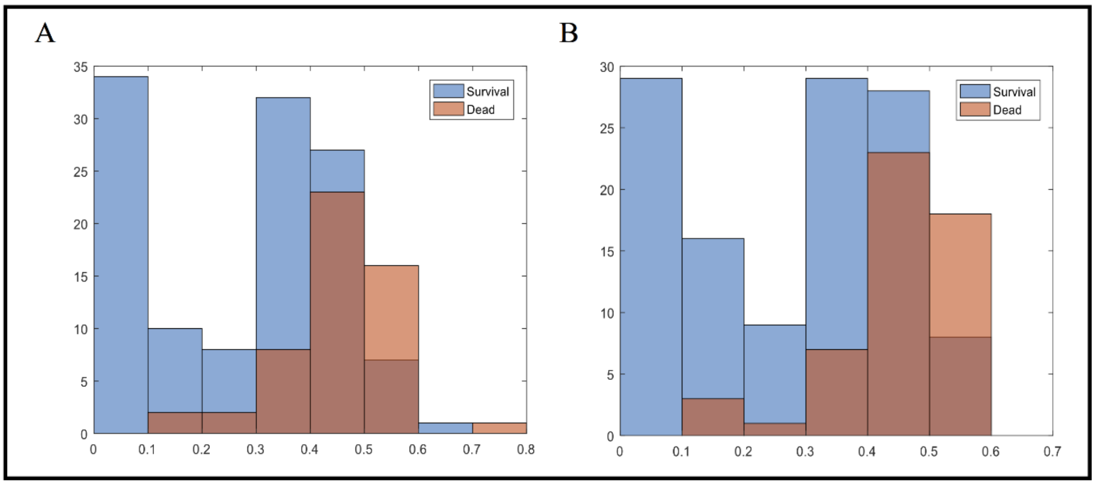


**Figure S2.** The probability distribution of patients in (A) microarray DNN and (B) integrative DNN in the test set.

We also found similar results for the independent validation set. We observed that two groups of probabilities (survival and death) were still not eminent in the microarray DNN. Specifically, it did not classify the [Probability < Youden index] region well in the microarray DNN. In other words, the integrative DNN was more generalizable after including the microarray and clinical data.

**Table S13: The distribution of patients in three regions for microarray DNN in the independent validation set.**

| Probability region | Survival | Death |
|---:|---:|---:|
| Probability < Youden index | 29 | 15 |
| Youden index < Probability < 0.5 | 11 | 5 |
| 0.5 < Probability | 11 | 19 |

**Table S14: The distribution of patients in three regions for integrative DNN in the independent validation set.**

| Probability region | Survival | Death |
|---:|---:|---:|
| Probability < Youden index | 22 | 6 |
| Youden index < Probability < 0.5 | 18 | 11 |
| 0.5 < Probability | 11 | 22 |

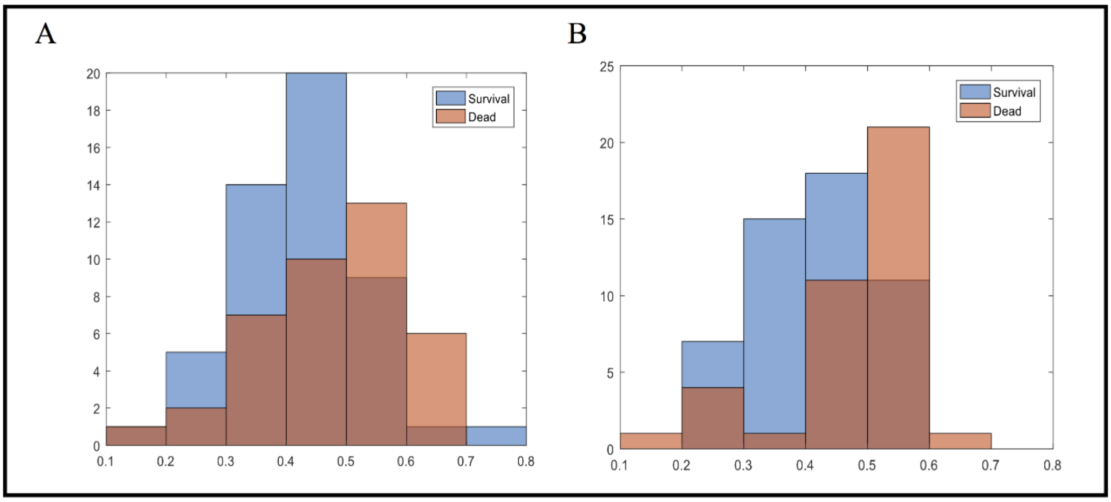


**Figure S3.** The probability distribution of patients in (A) the microarray DNN and (B) integrative DNN in the independent validation set.
